# Cumulus cell acetyl-CoA metabolism from acetate is associated with maternal age but inconclusively with oocyte maturity

**DOI:** 10.1101/2020.02.28.970327

**Authors:** Sharon Anderson, Peining Xu, Alexander J. Frey, Jason R. Goodspeed, Mary T. Doan, John J. Orris, Nicolle Clements, Michael J. Glassner, Nathaniel W. Snyder

**Affiliations:** Main Line Fertility, 825 Old Lancaster Road, Suite 170, Bryn Mawr, PA 19010; Ob/Gyn Department, Drexel University College of Medicine, Department of Obstetrics and Gynecology, Drexel University College of Medicine, Philadelphia, PA, USA; AJ Drexel Autism Institute, Drexel University, 3020 Market St Suite 560, Philadelphia, PA 19104; Center for Metabolic Disease Research, Department of Microbiology and Immunology, Temple University Lewis Katz School of Medicine. Philadelphia, PA, USA; Department of Decision System Sciences, St. Joes University, 348 Mandeville Hall, Philadelphia, PA, USA

**Keywords:** Cumulus cells, oocyte, metabolism, acetate metabolism

## Abstract

Cumulus cell (CC) clumps that associate with oocytes provide the oocytes with growth and signaling factors. Thus, the metabolism of the CCs may influence oocyte function and CC metabolism may be predictive of oocyte competence for in vitro fertilization. CCs are thought to be highly glycolytic but data on other potential carbon substrates are lacking in humans. This was a prospective and blinded cohort study that was designed to examine the substrate utilization of CCs by age and oocyte competence. Individual sets of CC clumps from participants were removed after oocyte retrieval procedure, incubated with stable isotope labeled substrates, and analyzed using liquid chromatography-high resolution mass spectrometry (LC-HRMS) for isotopologue enrichment of major metabolic intermediates, including acetyl-CoA. The acyl-chain of acetyl-CoA contains 2 carbons that can be derived from ^13^C-labeled substrates resulting in a M+2 isotopologue that contains 2 ^13^C atoms. Comparing the fate of three major carbon sources, mean enrichment of M+2 acetyl-CoA (mean, standard deviation) was for glucose (3.6, 7.7), for glutamine (9.4, 6.2), and for acetate (20.7, 13.9). Due to this unexpected high and variable labeling from acetate, we then examined acetyl-CoA mean % enrichment from acetate of in 278 CCs from 21 women ≤34 (49.06, 12.73) decreased with age compared to 124 CCs from 10 women >34 (43.48, 16.20) (p=0.0004, t test). The CCs associated with the immature prophase I oocytes had significantly lower enrichment in M+2 acetyl CoA compared to the CCs associated with the metaphase I and metaphase II oocytes (difference: −6.02, CI: −1.74,-13.79, p=0.013). Acetate metabolism in individual CC clumps was positively correlated with oocyte maturity and decreased with maternal age. These findings indicate that CC metabolism of non-glucose substrates should be investigated relative to oocyte function and age-related fertility.

## Introduction

A major barrier in in vitro fertilization (IVF) remains that approximately 5% of aspirated human oocytes have the competence after fertilization to implant and develop into a child [1]. Oocyte developmental competence decreases as a woman ages, and currently maternal age is the single best predictor of reproductive outcomes in women, highlighting an acute need for non-invasive biomarkers of oocyte competency to increase efficiency, lower costs, and reduce the chances of riskier multiple births [2]. This has led to attempts to understand the aging and maturation process of oocytes for both etiologic understanding of fertility and predictive assays for oocyte function and oocyte competence.

Notably, the metabolism of the oocyte in vivo is intimately linked to surrounding cells, such as the oocyte-associated somatic cumulus cells (CCs), which are critical to oocyte function in the ovarian follicle [3]. CCs support oocyte development through the provision of essential nutrients and signaling molecules [4]. CCs possess specialized cytoplasmic projections that penetrate through the zona pellucida [shell] and form gap junctions at their tips with the oocyte, generating an elaborate structure called the cumulus-oocyte complex (COC) [5, 6]. Within the COC, CCs are thought to metabolize the bulk of the glucose to supply metabolic intermediates to the oocyte [7], where the COC glucose metabolism is pivotal in determining oocyte developmental competence [8]. Because of the metabolic and communication link between the cumulus and the oocyte, glucose availability and metabolism within the cumulus can have a significant impact on oocyte meiotic and developmental competence [9].

During the oocyte retrieval procedure after controlled ovarian stimulation for IVF, the oocytes within specialized ovarian follicles, that support the growth and development of oocytes, are harvested. As only the oocyte is used in fertilization, the remainder of the cells and fluid (CCs, follicular fluid and ganulosa) is remnant sample and can be used as a source for non-invasive investigation of follicle function, even by destructive methods that consume the sample completely. This circumvents the issue of applying such methods to the oocyte which must be fertilized and implanted. This easy access has led to insightful investigations into the metabolomics of the follicular fluid [10], but relatively few studies have been conducted on the metabolism of the recovered human CCs.

In this study we utilized stable isotope tracing coupled with liquid chromatography-mass spectrometry (LC-MS) to quantify the substrate preferences for CC metabolism. This approach for ex vivo metabolic studies uses incubation with a stable (non-radioactive) isotope labeled metabolic substrate (e.g. ^13^C_6_-glucose) and tracking the incorporation of that isotopic label into downstream metabolites. With detection by LC-MS, the isotopologues (molecules differing only in their number of isotopes), co-elute on the LC, but are resolved by the MS, allowing separate quantitation of each isotopologue. After performing an adjustment for natural isotope abundance, the resulting isotopologue enrichment gives the relative molar % of the product detected derived from the labeled substrate in the given incubation conditions. Importantly, this isotopologue enrichment is self-normalizing and is not as influenced by the number of CCs in each sample which is variable.

Based on previous literature that CCs are highly glycolytic, we expected to see utilization of glucose to generate anabolic intermediates, including the central metabolic intermediate acetyl-Coenzyme A (acetyl-CoA). Acetyl-CoA is the acyl-donor for protein and histone acetylation, the source of two-carbon units for de novo fatty acid and cholesterol synthesis, and a source of two-carbon units for oxidation in the TCA cycle [11]. However, initial experiments demonstrated that acetyl-CoA was poorly labeled by glucose, and that acetate, not glucose or glutamine, was the preferred substrate to generate the central metabolic intermediate acetyl-CoA in the recovered CCs. Little is known about the metabolism of short chain fatty acids like acetate in the CCs, despite the importance of acetate in the contexts of neuronal metabolism, liver metabolism, and cancer [11, 12]. Thus, based on the finding of relatively high acetate usage to generate acetyl-CoA, we examined the association between cumulus cell acetate metabolism to acetyl-CoA, maternal age, and oocyte maturity.

## Results

### Cumulus cells have a high capacity to derive acetyl-CoA from acetate relative to glucose and glutamine

Cumulus cells incubated with either [^13^C_6_]-glucose, [^13^C_5_]-glutamine, or [^13^C_2_]-acetate for one hour were analyzed for isotopologue enrichment of the central carbon intermediate acetyl-CoA. Isotopologue enrichment, calculated with FluxFix (10), gives the relative molar % of the product detected derived from the labeled substrate in the given incubation conditions. Thus, in this context, the isotopologue notation describes how many labeled carbons are in the analyte of interest corresponding to M+0 as the unlabeled acetyl-CoA, M+1 containing one ^13^C, M+2 containing two ^13^C atoms (Figure 1). M+2 represents the maximum labeling of the acyl-group (the acetyl group) in acetyl-CoA. Mean isotopologue enrichment of M+2 acetyl-CoA (mean, standard deviation) was for glucose (3.6, 7.7), for glutamine (9.4, 6.2), and for acetate (20.7, 13.9) (Figure 2).

**Figure 1.**
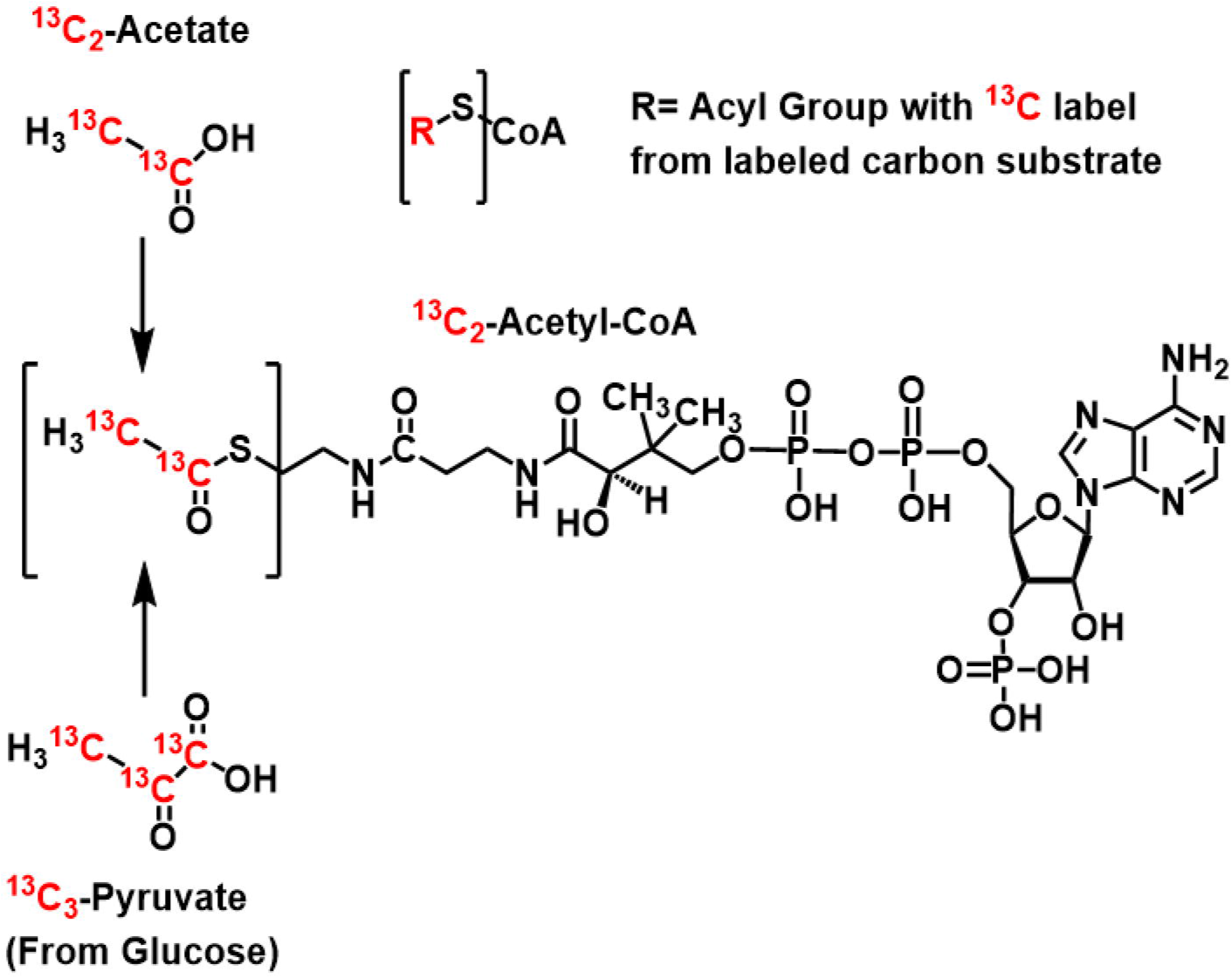
Stable isotope metabolic tracing can determine the substrates that contribute to the central metabolite Acetyl-CoA. Metabolic tracing from (A) Generally, acyl-CoA labeling can occur by any metabolite where the R=acyl group can be variably labeled by carbon derived from substrates. Stable isotope labeling into ^13^C_2_-acetyl-CoA can be achieved through ^13^C_2_-acetate or ^13^C_3_-pyruvate derived from ^13^C_6_-glucose.

**Figure 2.**
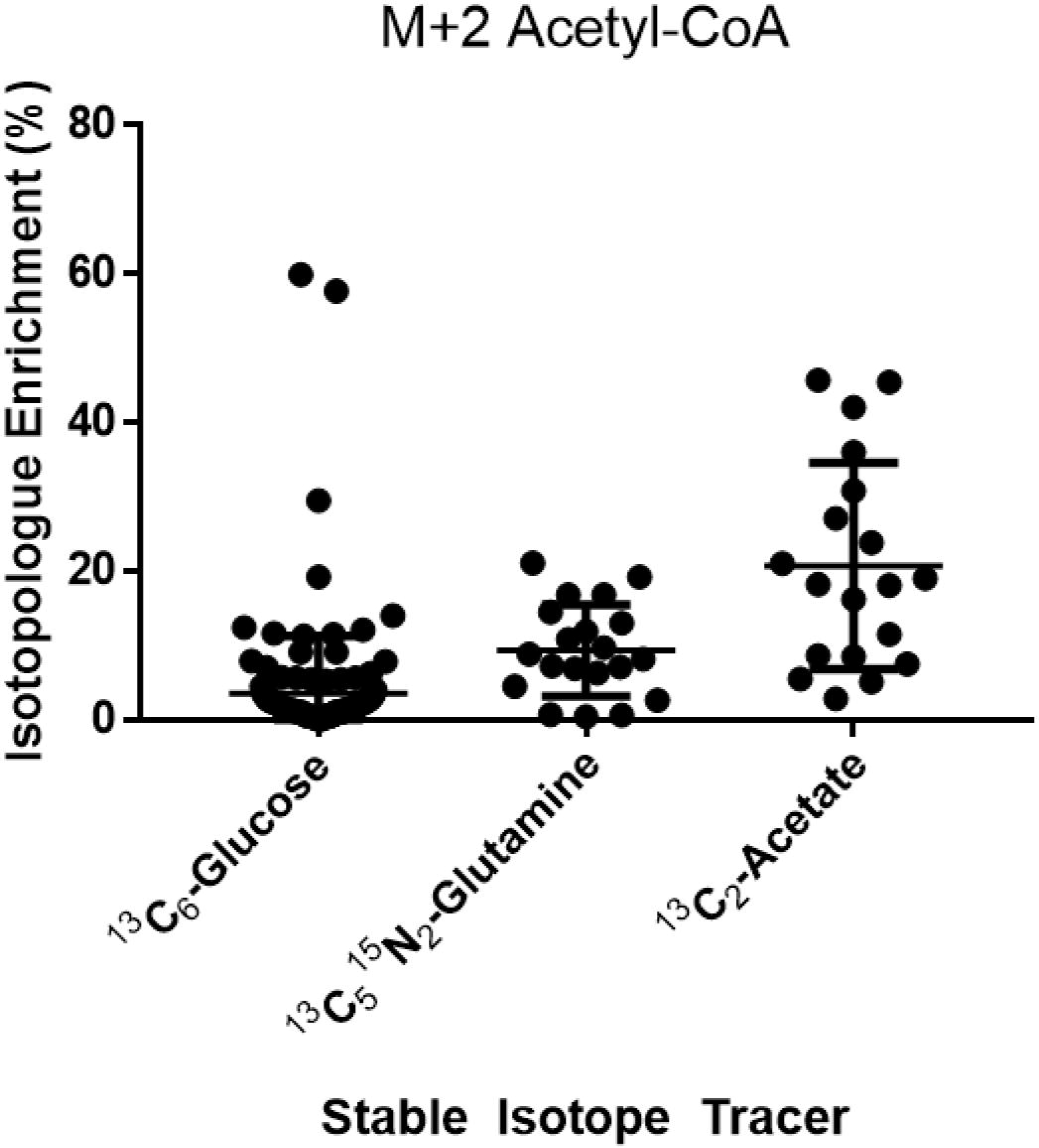
Isotopologue enrichment in acetyl-CoA M+2 from 3 major carbon tracers in cumulus cells (CCs) indicates that acetate can contribute significantly to acetyl-CoA. One hour incubations with CC clumps from individually retrieved oocytes in stable isotope tracer media containing the indicated labeled substrate. CCs have a higher capacity for generating acetyl-CoA from acetate than glucose, despite the literature on the glycolytic nature of glucose.

### Enrichment of acetyl-CoA from acetate is decreased in cumulus cells from older women

Both measures of metabolism and fertility are known to decline with age. To examine if this was true in CCs, we tested if acetyl-CoA enrichment from acetate was different from women ≤34 than >34. This cut-off is based on the distribution of ages within our sample, the age categories reported by the Society for Assisted Reproduction (SART), and the American College of Obstetricians and Gynecologists (ACOG) recognition of age 35 as a guideline for increased fertility counseling. Mean % enrichment of acetyl-CoA in 278 CCs from 21 women ≤34 (49.06, 12.73) was higher (p=0.0004, t test) than in 124 CCs from 10 women >34 (43.48, 16.20) (Figure 3).

**Figure 3.**
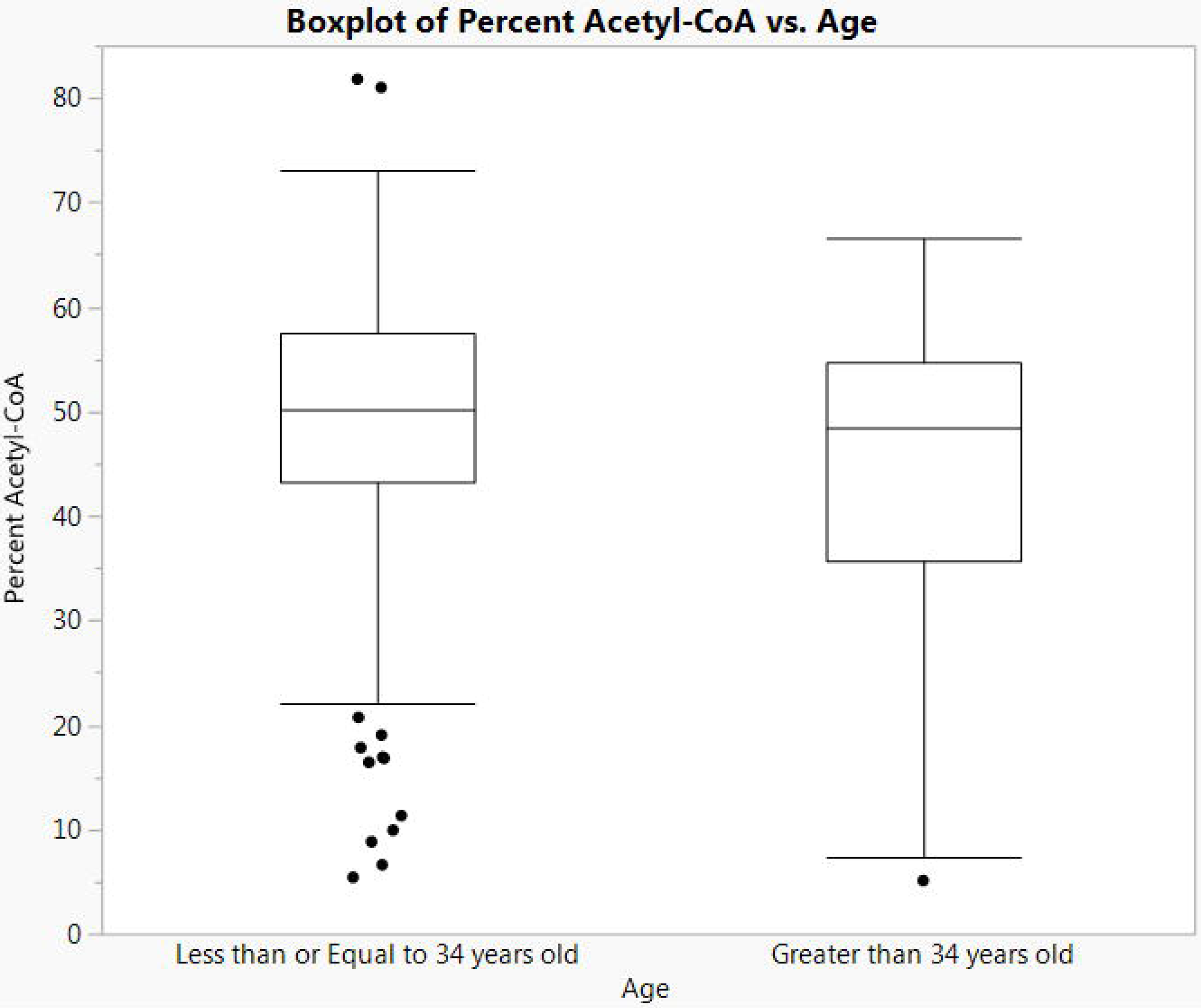
Isotopologue enrichment in acetyl-CoA M+2 in cumulus cells (CCs) varies by age group. Isotopologue enrichment in acetyl-CoA CCs from ^13^C_2_-acetate was 4.3% higher in women ≤34 (n=21) versus >34 (n=10) (p= 0.0004, CI: 2.3, 8.8).

Although we had pre-specified 34 as a cutoff before statistical analysis, we conducted an exploratory analysis to examine the effect of treating age as a continuous variable. The equation of the regression was estimated at % acetyl-CoA = 54.46 – 0.215(age), with a p-value for the slope coefficient of 0.314. The high variance in acetyl-CoA labeling and the restriction of our sample population to ages 26-42 limit the utility of this analysis.

### Generation of acetyl-CoA from acetate is associated with mature oocytes

Based on the discovery that cumulus cells have a high capacity to utilize acetate to generate acetyl-CoA, we were interested if the enrichment in acetyl-CoA from acetate correlated with properties of the associated oocyte, especially oocyte maturity. For CCs from 31 patients where oocyte maturity data was available, we tested if M+2 acetyl-CoA enrichment from acetate was different in cumulus cells associated with oocyte maturity. Three types of oocytes are obtained at retrieval – mature metaphase II oocytes, immature metaphase I oocytes (no polar body, and very immature prophase I oocytes (no polar body, contain a germinal vesicle) The cumulus cells associated with the immature prophase I group had significantly lower enrichment in M+2 acetyl CoA compared to the cumulus associated with the other oocytes (difference: 7.77, CI 1.74, 13.79, p=0.013) (Figure 4).

**Figure 4.**
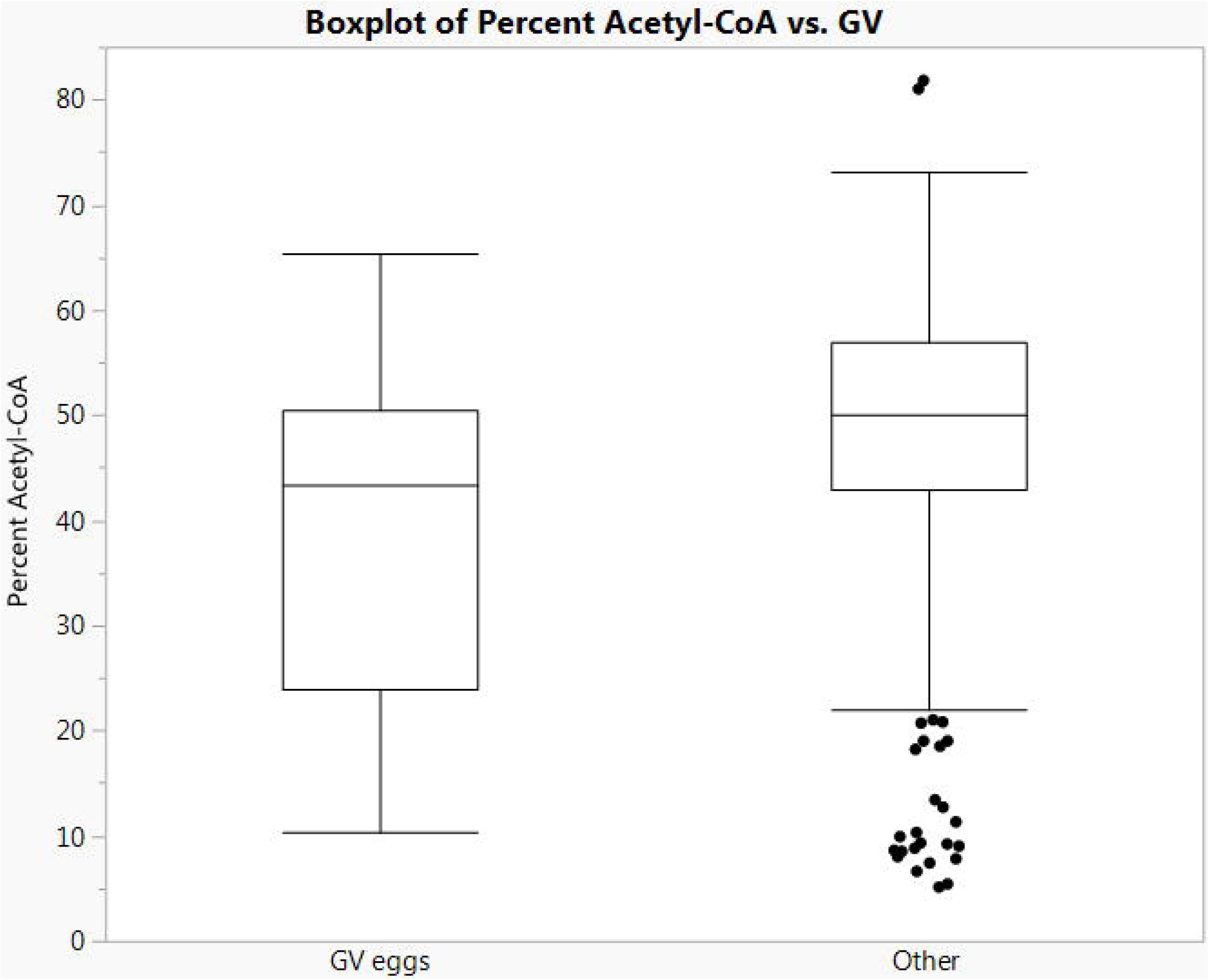
Isotopologue enrichment from CCs associated with prophase I oocytes containing germinal vesicles (GV eggs) is lower than that from CCs associated with metaphase I and II oocytes. Isotopologue enrichment in acetyl-CoA CCs associated with GV eggs was 6.02% lower than in CCs associated with metaphase I and II oocytes (other) (p=0.013, CI: −1.74,−13.79).

Finally, we examined if there was a difference in acetyl-CoA isotopologue enrichment between CCs that were associated with embryos that were selected for transfer to the uterus versus those that were not. CCs that were associated with embryos that were not selected for transfer had no significant difference in isotopologue enrichment (−2.54, CI: 3.29, −8.38, p=0.38). Similarly, comparing SART scoring of embryo quality (Grade A vs other grades) on day 3 of embryo culture, there was no significant difference in isotopologue enrichment (1.46, CI: 4.9 to −1.97, p=0.40). Comparing differences by all embryo grades, there were no significant differences in acetyl-CoA isotopologue enrichment between groups (p=0.51).

## Discussion

Our findings indicate that cumulus cells have a high capacity to derive acetyl-CoA from acetate. This also partially contrasts to previous literature, implicating cumulus cells as highly glycolytic, since labeling was conducted in the presence of 5mM unlabeled glucose for glutamine and acetate tracing. Importantly, this suggests that the role of glycolysis in CCs may not be to feed acetyl-CoA, via conversion to pyruvate, into the TCA cycle for oxidation or export as citrate to generate cytoplasmic acetyl-CoA. This alternative use of glucose would be in-line with reports that CCs use glucose to generate extra-cellular matrix components or increased products of the hexosamine pathway [13]. Future studies will need to test the effect of acetate concentration closer to reported physiological circulating blood levels of 50-100 μM acetate. Additionally, the use of other tracers including pyruvate, and the measurement of isotope incorporation into other analytes including TCA cycle intermediates including malate and citrate that are involved in transport of carbon from the mitochondria to the cytosol, could identify the specific enzymes responsible for our observations.

Strengths of this study include the novelty of applying stable isotope assisted metabolomics to individual oocyte-associate cumulus cell complex metabolism. Studying the ex vivo metabolism of human cumulus cells is difficult because of the paucity of recovered cells associated with single oocytes, variability in the numbers of cells, and the effects of endogenous metabolism beyond the control of the experimentalist. With stable isotope tracing, it is possible to establish a substrate/product relationship, in comparison to the metabolomics experiments already conducted that only examined the contents of the follicular fluid with no regard as to their origin. Isotope tracing performed ex vivo reduces the major confounder of diet and resulting changes in circulating metabolites. Additionally, isotope tracing is “self-normalizing” such that variation in cell numbers are accounted for because each isotopologue is normalized to the metabolite pool within each sample (e.g. acetyl-CoA M+2 is a fraction of total acetyl-CoA in that sample) [14]. Finally, since this study used human cumulus cells, our interpretation of the role of short chain fatty acids including acetate may be distinct from that in ruminants or rodents where higher levels of acetate, propionate, and butyrate are found in circulation.

Although we found a difference in acetyl-CoA labeling by oocyte maturity, a limitation of this study is that we did not follow through data collection to oocyte competence to pregnancy or to baby. Oxygen consumption, a rough gauge of metabolic activity, is an existing quality marker for human oocyte competence conditioned by ovarian stimulation regimens [15]. Furthermore, the oxygen consumption rates of embryos have been found to be associated with successful implantation and can be used to select the embryo with the best developmental potential [16]. However, this quantitation of metabolic function was a relatively poor predictor, with an odds ratio of 1.037-1.732 (95% CI) for normal vs non-fertilized/abnormal oocytes [15]. Future work may be able to improve upon this prediction incorporating more specific substrates in combination with oxygen consumption. The finding of high levels of acetate utilization, and significant glutamine labeling into acetyl-CoA suggests that exploration of other potential substrates may have utility. To this end, use of ketone body (beta-hydroxybutyrate) and fatty acid tracers may be particularly insightful in future studies.

Our finding of decreased acetyl-CoA generation from acetate in cumulus cells from women >35 is biochemically interesting. The aging process of the oocyte is complex and includes inter-related impaired mitochondrial dysfunction, oxidative stress, diminished metabolic activity, potential accumulation of DNA damage, and activity of several cell-signaling systems [17]. Pathological perturbations of the coordinated somatic cell-oocyte interactions by metabolic disease and/or maternal aging can induce molecular damage to the oocyte can alter macromolecules, induce mitochondrial mutations, all of which can harm the oocyte [18]. It is also thought that these pathological processes harm the related CCs. Endometriosis may be associated with mitochondrial dysfunction in pooled CCs [19], and subjects with endometriosis may have a defect in CC mitochondrial function, which likely contribute to decreased fertilization and implantation rates [20]. Generation of acetyl-CoA is possible from a variety of substrates, but two major sources include glucose and acetate. A switch between glucose and acetate is alternatingly dependent on the enzymes ATP citrate lyase (ACLY) that generates glucose-derived acetyl-CoA from citrate exported from the mitochondria and the Acyl-coenzyme A synthetase short-chain family members that form acetyl-CoA from acetate and CoASH [12]. In this study, our exclusion criteria limits the extrapolation of our findings to wider age ranges and pathological settings. Future studies using stable isotope labeling of cumulus cells may be useful in non-invasive study of the metabolic consequences of endometriosis and other diseases, especially in the context of fertility.

## Materials and Methods

### Chemicals

Water, methanol, acetonitrile, and ammonium acetate were Optima LC-MS grade solvents from Fisher Scientific (Pittsburgh, PA). Trichloroacetic acid and salts used in Tyrode’s buffer were from Sigma-Aldrich (St. Louis, MO). Stable isotope-labeled substrates were from Cambridge Isotope Laboratories (Tewksbury, MA).

### Patient Selection

Enrolled patients were scheduled for IVF at the Main Line Fertility Center (Bryn Mawr, Pennsylvania, USA). Women undergoing IVF were screened for inclusion in this study and informed consent was obtained from all individual participants before enrollment. Inclusion criteria included IVF patients between the ages of 28 and 42 years, with an anti-mullerian hormone (AMH) between 1.0 and 10 ng/ml, and follicle stimulating hormone ≤ 10 IU/ml, luteinizing hormone (LH) < 12 IU/ml, and estradiol < 50 pg/ml on day 2-4 of the menstrual cycle. A body weight ≥ 50 kg and a body mass index between 18 and 32 kg/m^2^ were required. Patients were excluded for smoking, polycystic ovarian disease, endometriosis greater than Stage I, utilizing testicular sperm for IVF, and preimplantation genetic testing.

### Controlled Ovarian Stimulation Protocol

Patients underwent the standardized controlled ovarian stimulation for IVF using Gonal-F RFF Redi-ject pen (EMD Serono, Inc., Darmstadt, Germany). All patients received a fixed protocol of 300 IU recombinant FSH (rFSH) daily for the first four days of stimulation. After, a flexible protocol was used to optimize ovarian response. rFSH was adjusted by the patient’s physician to 225 to 450 IU daily up to and including day of human chorionic gonadotropin (hCG) trigger administration. Cycles were monitored with follicular ultrasound measurements and serum estradiol concentrations throughout controlled ovarian stimulation. An antagonist was used to suppress endogenous pituitary LH for the prevention of premature LH surges by administering 0.25 mg/day of Cetroelix Acetate (EMD Serono) when follicle size reached 12 mm and continued up to and including day of hCG trigger. Subcutaneous injection of 250 μg of hCG (Ovidrel, EMD Serono) was administered when at least three leading follicle sizes reached a diameter of ≥ 17 mm. The oocyte retrieval procedure was performed 36 h after hCG injection.

### Biosample Collection

CCs were removed from cumulus-oocyte complexes immediately after the oocyte retrieval procedure as part of routine IVF lab protocol. Instead of being discarded, CCs were placed into individually labeled micro-centrifuge tubes containing one mL of transport medium (HEPES ((4-(2-hydroxyethyl)-1-piperazineethanesulfonic acid)-buffered medium containing 5 mg/ml human serum albumin from CooperSurgical (Catalog ART-1012, Trumball, CT), then delivered by medical courier with a 37°C heat pack to a university research laboratory where metabolic tracing and analysis were conducted. 31 patients and a total of 402 CC clumps had paired with oocyte maturity data with CCs from each stage as follows; 0PN, 34; 1PN, 8; 2PN, 233;3PN, 6;A, 11;GV, 30;MI, 44;MII, 32. This removal was before preparation for other procedures, thus the CCs used for this study were not exposed to hyaluronidase.

### Cumulus Cell Metabolic Tracing

All tracing was conducted by laboratory analysts blinded to any sample characteristics using adaptations of previously developed methods [21]. CC metabolism was traced by centrifugation of the CC clump at 200 × g for 5 min at 25°C, removal of the supernatant, and then addition of the labeling media. For labeling media we used Tyrode’s buffer (139 mmol/L NaCl, 3 mmol/L KCl, 17 mmol/L NaHCO_3_, 3 mmol/L CaCl_2_, and 1 mmol/L MgCl_2_) pH= 7.4 pre-incubated at 20% O_2_ and 5% CO_2_ containing as a carbon source either 5 mmol/L [^13^C_6_]-glucose or 5 mmol/L glucose in combination with either 1 mmol/L [^13^C_2_]-acetate or 100 μmol/L [^13^C_5_ ^15^N_2_]-glutamine. Labeling was conducted by addition of labeling media, a 3 sec vortex, and then incubation in a 37°C water bath for 1 hour in 1.5 mL plastic Eppendorf tubes. After completion of labeling incubation, cumulus cells were centrifuged at 1000 × g for 5 min at 4°C, supernatant was removed, and either 1 mL of 80:20 methanol:water pre-chilled to −80°C (for lactate and pyruvate analysis) or 1 mL of 10% trichloroacetic acid (w/v) in water (for acyl-CoA analysis) was added. Samples were frozen at −80°C until extraction and analysis. Samples were extracted by thawing at 4°C, probe tip sonication for 15 sec, and then centrifugation at 16,000 × g for 10 min at 4°C. Supernatant was extracted with an Oasis HLB solid phase extraction (SPE) cartridge washed with 1 mL methanol and then equilibrated with 1 mL water. After loading of the supernatant, the SPE column was washed with 1 mL water, then eluted into a 10 mL glass tube with 1 mL methanol with 25mM ammonium acetate. The eluent was evaporated to dryness under nitrogen, resuspended in 50 μL 5% (w/v) 5-sulfosalicylic acid and 10 μL was injected for LC-HRMS analysis.

### Liquid Chromatography-High Resolution Mass Spectrometry (LC-HRMS)

Acyl-CoAs were analyzed as previously described in a quantitatively validated method [22] on an Ultimate 3000 Quaternary UHPLC coupled to a Q Exactive Plus mass spectrometer operating in the positive ion mode with a heated electrospray ionization probe (mark II) in an IonMax Source housing. Samples were kept in a temperature controlled autosampler at 6°C and LC separation was performed as previously described on a Waters HSS T3 2.7 μm particle size 2.1 × 150 mm column. LC conditions were as follows; column oven temperature 30°C, solvent A water with 5 mM ammonium acetate, solvent B 95:5 acetonitrile: water with 5 mM ammonium acetate, solvent C (wash solvent) 80:20 acetonitrile: water with 0.1% formic acid. The gradient was as follows: 0.2 mL/min flow at 98% A and 2% B for 1.5 min, 80% A 20% B at 5 min, 100% B at 12 min, 0.3 mL/min 100% B at 16 min, 0.2 mL/min 100% C at 17 min, held to 21 min, then re-equilibrated at 0.2 mL/min flow at 98% A and 2% B from 22 to 28 min. Flow from 4-18 minutes was diverted to the instrument. Operating conditions on the mass spectrometer were as follows; auxiliary gas 10 arbitrary units (arb), sheath gas 35 arb, sweep gas 2 arb, spray voltage 4.5 kV, capillary temperature 425°C, S-lens RF-level 50, aux gas heater temperature 400°C, in-source CID 5 eV. Scan parameters were optimized during the experiments, but final conditions were alternating full scan from 760-1800 *m/z* at 140,000 resolution and data independent acquisition (DIA) looped 3 times with all fragment ions multiplexed at a normalized collision energy (NCE) of 20 at a resolution of 280,000. An isolation width of 7 *m/z* with an offset of 3 *m/z* was used to capture all relevant isotopologues for targeted acyl-CoAs. Data was processed in Xcalibur and or TraceFinder (Thermo).

## Data analysis

Normalization of isotopologue distribution by natural isotopic abundance was conducted via an open source resource, FluxFix [14], using experimentally derived normal isotopic distributions from cumulus cell clumps labeled with non-isotopically enriched glucose. Due to this adjustment, any sample that had zero signal intensity for M+0 or M+2 was excluded from analysis, since these values would not be interpretable. For isotopologue analysis, summary statistics were calculated in Graph Pad Prism (v7).

Metabolic analysis was conducted by investigators blinded to status and identity of all samples/patients, and analysis of correlation with oocyte maturity and age was conducted by an independent biostatistician, who had no role in the design or collection of data, using JMP Pro 13.0 software. Unit of analysis was on each cumulus cell clump sample, since we were primarily interested in the unique metabolism of the cumulus cells associated with individual oocytes.

## Ethics approval

Study protocol was approved by Western Institutional Review Board (WIRB) study number 1160207 under investigator Glassner and all patients were consented to participate in this research.

## Author Contribution

Conducted experiments and analyzed data (PX, AJF, JRG, and MTD). Analyzed data (NC). Conducted experiments and obtained samples (SA, JJO, and MJG). Wrote the manuscript, obtained funding, conducted experiments, and analyzed data (SA and NWS).

## Funding/Acknowledgements

Funding for this research was provided in part by EMD Serono, Inc./EMD Serono Research & Development Institute, Inc. The funders had no role or input in design, conduct, analysis, interpretation or reporting of the research.

